# lncRNA *HOTAIRM1* Coordinates with RNA Processing Factors in DNA Damage Repair

**DOI:** 10.1101/2022.08.13.503833

**Authors:** Tzu-Wei Chuang, Pei-Yu Wu, Yao-Ming Chang, Woan-Yuh Tarn

**Author notes:** Corresponding author: Woan-Yuh Tarn, Ph.D., Institute of Biomedical Sciences, Academia Sinica, 128 Academy Road Section 2, Nankang, Taipei 11529, Taiwan.

## Abstract

The eukaryotic RNA processing factor Y14 participates in double-strand break (DSB) repair via its RNA-dependent interaction with the non-homologous end-joining (NHEJ) complex. We identified the long non-coding RNA *HOTAIRM1* as a candidate that mediates this interaction. *HOTAIRM1* localized to DNA damage sites induced by ionizing radiation. Depletion of *HOTAIRM1* delayed the recruitment of DNA damage response and repair factors to DNA lesions and reduced DNA repair efficiency. Identification of the *HOTAIRM1* interactome revealed a large set of RNA processing factors including mRNA surveillance factors. The surveillance factors Upf1 and SMG6 localized to DNA damage sites in a *HOTAIRM1*-dependent manner. Depletion of Upf1 or SMG6 increased the level of DSB-induced non-coding transcripts at damaged sites, indicating a pivotal role for Upf1/SMG6-mediated RNA degradation in DNA repair. We conclude that *HOTAIRM1* serves as an assembly scaffold for both DNA repair and RNA processing factors that act in concert to repair DSBs.

## INTRODUCTION

DNA damage may arise from physiological processes or can be generated by environmental toxins and/or reactive oxygen species (Jackson and Bartek, 2009). Inefficient repair of DNA lesions threatens genome stability and underlies a number of human diseases, particularly cancer. The diversity of DNA lesion types necessitates multiple distinct DNA repair mechanisms. DNA double-strand breaks (DSBs) are generated by ionizing radiation (IR), severe oxidative damage, transcriptional R-loops, or replication fork collapse (Lieber, 2010). DNA damage triggers the DNA-damage response (DDR), which comprises a network of cellular pathways that sense and repair DNA lesions. Moreover, DDR can activate cell-cycle checkpoints to safeguard genome integrity or cause apoptosis (Giglia-Mari et al., 2011). DSBs represent one of the most hazardous lesions. The Mre11-Rad50-Nbs1 (MRN) complex acts as a DSB sensor and coordinates DSB repair. Two principal mechanisms are used for DSB repair, namely nonhomologous end-joining (NHEJ) and homologous recombination (HR) (Lieber, 2010; Pannunzio et al., 2018). NHEJ can function throughout the cell cycle and hence is the dominant DSB repair pathway, albeit with a higher error rate. The core repair machinery of NHEJ comprises a set of proteins including the Ku70/80 heterodimer, DNA-dependent protein kinase (DNA-PK) catalytic subunit, the DNA endonuclease Artemis, DNA ligase IV, XRCC4-like factor (XLF), and DNA polymerases λ/µ. The binding of Ku70/80 to DSB ends activates DNA-PK and NHEJ (Lieber, 2010; Pannunzio et al., 2018).

In addition to canonical DNA repair factors, RNA processing factors are also critical for maintenance of genome stability. Some of these factors control the expression of DNA repair proteins or directly participate in DNA damage responses via interaction with DNA damage response factors and repair factors. For example, BCLAF1 cooperates with phosphorylated BRCA1 to regulate the splicing of the transcripts encoding DNA repair factors after DNA damage (Savage et al., 2014). SFPQ/PSF promotes HR via interactions with the homologous recombinase Rad51 and the topoisomerase-binding protein TopBP1, and likewise promotes NHEJ via interaction with XRCC4-like factor (NHEJ1/XLF) (Jaafar et al., 2017; Morozumi et al., 2009). Poly(ADP-ribose) polymerase 1 (PARP1) catalyzes the formation of poly(ADP-ribose) onto itself or target proteins immediately after DNA damage (Azarm and Smith, 2020). Certain RNA-binding proteins (RBPs), such as SAF-A/hnRNP U, EWS and FUS/TLS, are recruited to DNA damage sites in a poly(ADP-ribose)-dependent manner (Altmeyer et al., 2015; Britton et al., 2014; Rulten et al., 2014; Savage et al., 2014). Localized RBPs can form liquid phase-separated compartments that facilitate the recruitment of DSB repair factors (Kai, 2016; Klaric et al., 2021). However, exactly how the various RBPs function in DSB repair awaits investigation.

Besides RBPs, RNAs also play a role in DDR or DNA repair (Domingo-Prim et al., 2020; Hawley et al., 2017; Wu and Wang, 2017). Indeed, DNA damage-induced long non-coding RNAs (lncRNAs) can regulate gene expression via interactions with DNA/RNA-binding proteins such as p53, TLS and YBX1 (Diaz-Lagares et al., 2016; Schmitt et al., 2016; Wang et al., 2008). For example, *Norad* sequesters Pumilio proteins to suppress the expression of mitotic and DNA repair factors and may also form a complex with several RBPs and topoisomerase I to ensure proper cell-cycle progression and chromosome segregation (Lee et al., 2016b; Munschauer et al., 2018). Furthermore, accumulating evidence suggests the existence of several different types of ncRNAs that directly participate in DSB repair. DSB sites can bidirectionally generate damage-induced long ncRNAs (dilncRNAs) through transcription by RNA polymerase II (Michelini et al., 2017; Pessina et al., 2019). The MRN complex facilitates such DSB induced transcription by melting DNA ends (Sharma et al., 2015). These dilncRNAs are subsequently processed into small RNAs in a Dicer/Drosha-dependent or -independent manner, preferentially at repetitive regions or ribosomal DNA loci (Bonath et al., 2018; Francia et al., 2012; Michelini et al., 2017). RNA hybrids formed by these dilncRNAs and small RNAs serve as a signal for recruiting the DDR factors MDC1 and 53BP1 to DNA damage sites (Michelini et al., 2017). The dilncRNAs can also form hybrids with resected single-stranded DNA ends in cell-cycle phases S and G2 to recruit HR factors (D’Alessandro et al., 2018). Besides the transcripts generated from DSB sites, several lncRNAs have direct roles in DSB repair. *BGL3* and *DDSR1* regulate BRCA1 accumulation at DSBs (Hu et al., 2020; Sharma et al., 2015). Several other lncRNAs, such as *LINP1* and *SNHG12*, mediate the interaction between DNA-PK and Ku70/80 and hence participate in NHEJ (Haemmig et al., 2020; Zhang et al., 2016). The mechanisms underlying the function of individual RNAs in DNA damage repair require further investigation.

The RNA processing factor Y14 functions in mRNA localization in the *Drosophila* germline and acts as a core factor of the exon-junction complex (EJC) in higher eukaryotes, which provides a link between splicing and nonsense-mediated decay (NMD) of mRNAs (Kurosaki et al., 2019). Y14 also modulates alternative splicing of precursor mRNAs, particularly those involved in apoptosis and cell-cycle progression (Fukumura et al., 2016; Michelle et al., 2012). Accordingly, depletion of Y14 causes cell-cycle arrest and apoptosis (Fukumura et al., 2016; Lu et al., 2017; Michelle et al., 2012). Several lines of evidence indicate that Y14 is important for maintenance of genome or chromosome integrity (Chuang et al., 2019; Lu et al., 2017; Silver et al., 2010). We reported that Y14, but not other EJC factors, specifically interacts with the NHEJ and DDR factors(Chuang et al., 2019). Depletion of Y14 causes delayed recruitment of these factors to DSB sites and thus impairs DNA repair. The notion that Y14 interacts with NHEJ factors in an RNA-dependent manner suggests the involvement of RNA in Y14-mediated DNA repair. To test this hypothesis, we identified Y14-associated lncRNAs that participate in the NHEJ pathway and explored the mechanism underlying RNA-mediated DSB repair.

## RESULTS

### Y14 is Associated with Non-coding RNAs

We previously established that RNA mediates the interaction between Y14 and Ku70/80 (Chuang et al., 2019). Y14 is more abundant than eIF4A3, another EJC factor, in the chromatin-enriched fraction, and further accumulates on chromatin with Ku70/80 after DNA damage (Chuang et al., 2019). To identify putative Y14-associated RNAs, we overexpressed FLAG-Y14 in HEK293 cells and performed RNA immunoprecipitation (RIP) using anti-FLAG with the chromatin-enriched fraction. Co-precipitated RNAs were subjected to high-throughput RNA sequencing using an Illumina HiSeq platform (Figure 1A). RIP-coupled with RNA sequencing was performed in duplicate (STAR Methods), which detected 28,593 expressed genes [FPKM (fragments per kilobase of transcript per million reads) values of >0 in at least one sample] (Table S1). Analysis with the NOISeq R package generated 6175 differentially expressed genes representing transcripts that co-immunoprecipitated with Y14. Reactome pathway enrichment analysis revealed that these Y14 target mRNAs encode proteins that function in DNA repair, the cell cycle, and RNA metabolism as the top-ranked annotations (Figure 1B; Table S2), consistent with previous findings that Y14 regulates the expression or splicing of DNA repair or cell cycle-related factors (Chuang et al., 2019; Fukumura et al., 2016; Michelle et al., 2012). Y14-associated mRNAs had a significantly greater number of exons compared with non-associated mRNAs, supporting the established role for Y14 as an EJC component (Figure 1C, left and middle). Approximately 5% (349) of the Y14-assoicated transcripts were lncRNAs (Table S1). Unlike the co-immunoprecipitated mRNAs, there was no difference in exon number between Y14-associated and non-associated lncRNAs (Figure 1C, right). Among the identified lncRNAs, ∼30% have been annotated, including small nucleolar RNA host genes (*SNHG*s) and previously characterized lncRNAs and antisense RNAs, whereas ∼70% were of unknown identity (Figure 1D). Figure 1E shows annotated lncRNAs having FPKM values of >20; notably, a set of *SNHG* genes was the highest ranked. We arbitrarily selected three *SNHG* RNAs and six lncRNAs for verification (Figure 1E, indicated by dots). Immunoprecipitation followed by analysis with reverse transcription-PCR (RT-PCR) showed that FLAG-Y14 associated with all the selected lncRNAs but not β-actin mRNA (control), validating the specificity of the affinity selection (Figure 1F). Finally, we observed that both FLAG-tagged Y14 and eIF4A3 interacted with two *SNHG* transcripts but not *HOTAIRM1* or *DANCR* (Figure 1G). It is possible that Y14 and eIF4A3 participate in NMD of *SNHG* RNAs (Lykke-Andersen et al., 2014).

**Figure 1.**
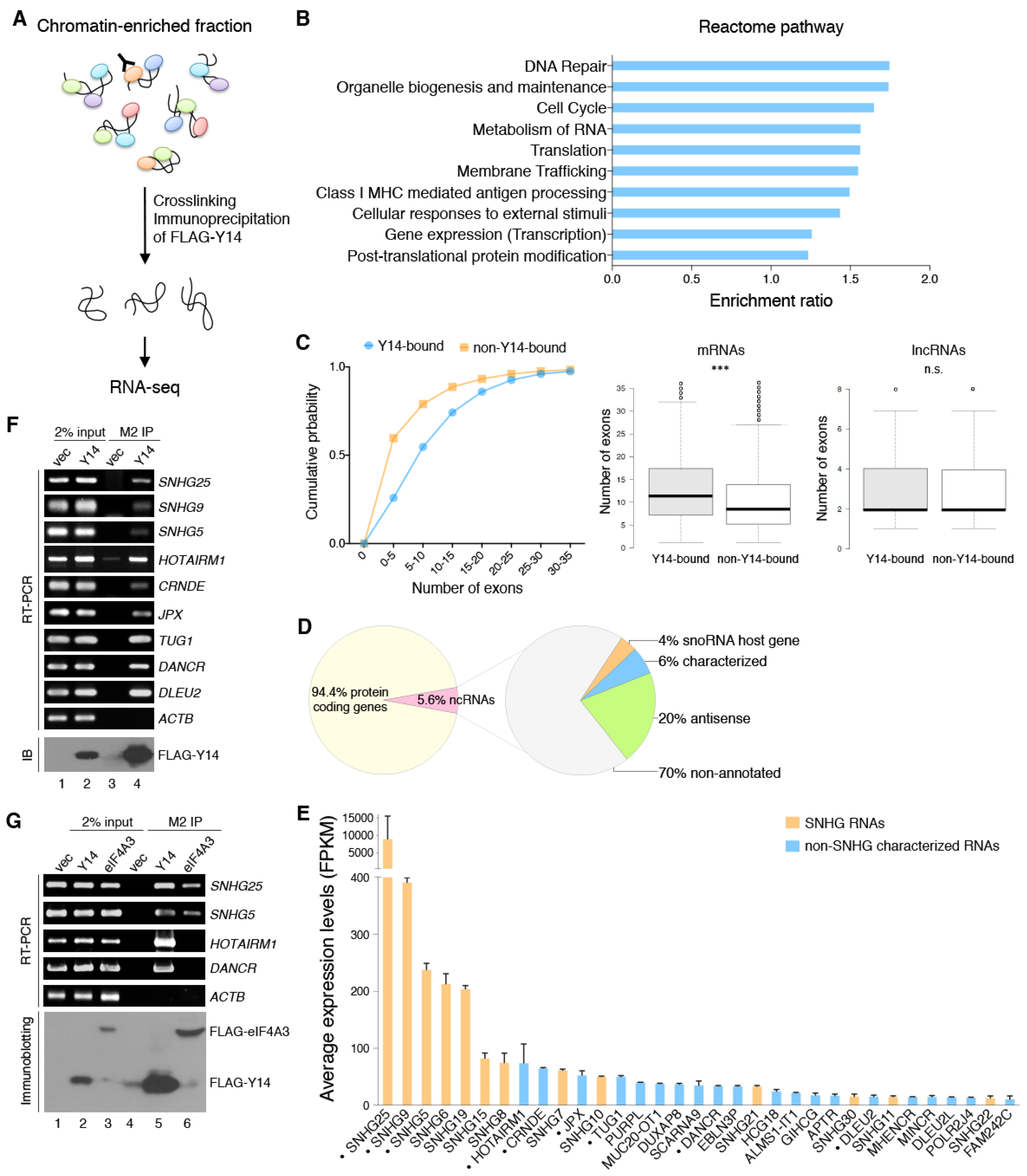
Identification of Y14-Associated RNAs from Chromatin-Enriched Fractions. (A) The diagram illustrates the procedure for identifying Y14-bound RNAs in the chromatin-enriched fraction of HEK293 cells that transiently expressed FLAG-Y14 through immunoprecipitation using anti-FLAG. (B) Reactome pathway analysis of proteins encoded by Y14-associated mRNAs. The bar graph shows the top 10 enriched pathways ranked by the enrichment ratio, *i*.*e*. the number of observed genes divided by the number of expected genes from each reactome pathway. (C) Graph showing the distribution of the number of exons per gene (Y14-bound RNAs vs. non-Y14-bound RNAs). The average number of exons of Y14-associated mRNAs (middle panel, \*\*\**p* < 0.001) and lncRNAs (right panel, n.s. not significant). (D) The percentage of different classes of Y14-associated RNAs (left, total 6175) and lncRNAs (right, total 349). Characterized lncRNAs indicate those with non-systematic symbols (genenames.org). snoRNA, small nucleolar RNA. (E) The 33 top annotated lncRNAs that were associated with Y14 (average from two independent RIP-Seq experiments). (F) HEK293 cells were transiently transfected with empty (vec) or FLAG-Y14-expressing vector. Cell lysates were subjected to immunoprecipitation (IP) using anti-FLAG, followed by RT-PCR using primers specific for the indicated lncRNAs or β-actin (*ACTB*, control). Immunoblotting was performed using anti-FLAG. (G) HEK293 cells were transiently transfected with empty (vec) or FLAG-Y14- or FLAG-eIF4A3-expressing vector. Immunoprecipitation, RT-PCR, and immunoblotting were as in panel F.

### *HOTAIRM1* Mediates the Interaction Between Y14 and Ku70/80

To pinpoint which lncRNA may have a role in the Y14-associated DNA repair pathway, we took advantage of two Y14 mutants, namely SA (S166A, S168A) and WV (W73V) (Chuang et al., 2016; Hsu Ia et al., 2005) (Figure 2A). As compared with wild-type Y14, the non-phosphorylatable Y14-SA was more abundant in the chromatin fraction (Figure S1A) and co-precipitated with Ku70/80 to a greater extent (Figure 2B, cp. lane 7 to lane 6). Y14-WV lacks RNA-binding activity (Chuang et al., 2016) and was unable to interact with Ku70/80 (Figure 2B, lane 8). Therefore, Y14 and its mutants exhibited differential binding affinity towards Ku70/80. Moreover, the observation that Y14-WV was unable to bind *HOTAIRM1* supported the idea that the Y14-Ku70/80 interaction is RNA dependent. Among the three lncRNAs noted above, only *HOTAIRM1* showed greater affinity for Y14-SA and no interaction with Y14-WV (Figure 2B), supporting its role as a mediator between Y14 and Ku. Indeed, *HOTAIRM1* co-precipitated with FLAG-Ku70 (Figure 2C). Moreover, using recombinant His-tagged Y14 and Ku70/80, we observed that both could pull down *HOTAIRM1* from HeLa cell total RNA (Figure 2D, lanes 4 and 5). However, His-tagged Y14-ΔC (lacking the C-terminal arginine-rich region) failed to pull down *HOTAIRM1* (Figure 2D, lane 3), indicating that this region is essential for the interaction of Y14 with *HOTAIRM1*. To examine whether the formation of the Y14-Ku70/80 complex depends on *HOTAIRM1*, we immunoprecipitated FLAG-Y14 from HEK293 cell lysates followed by RNase H digestion in the presence of an oligonucleotide complementary to *HOTAIRM1* (Figures 2E and S1B, AS1 and AS2). Two antisense *HOTAIRM1* oligonucleotides substantially disrupted the interaction between FLAG-Y14 and Ku70/80, whereas the nonspecific or antisense ACTB oligonucleotide had no effect (Figure 2F), supporting a scaffold role for *HOTAIRM1*.

**Figure 2.**
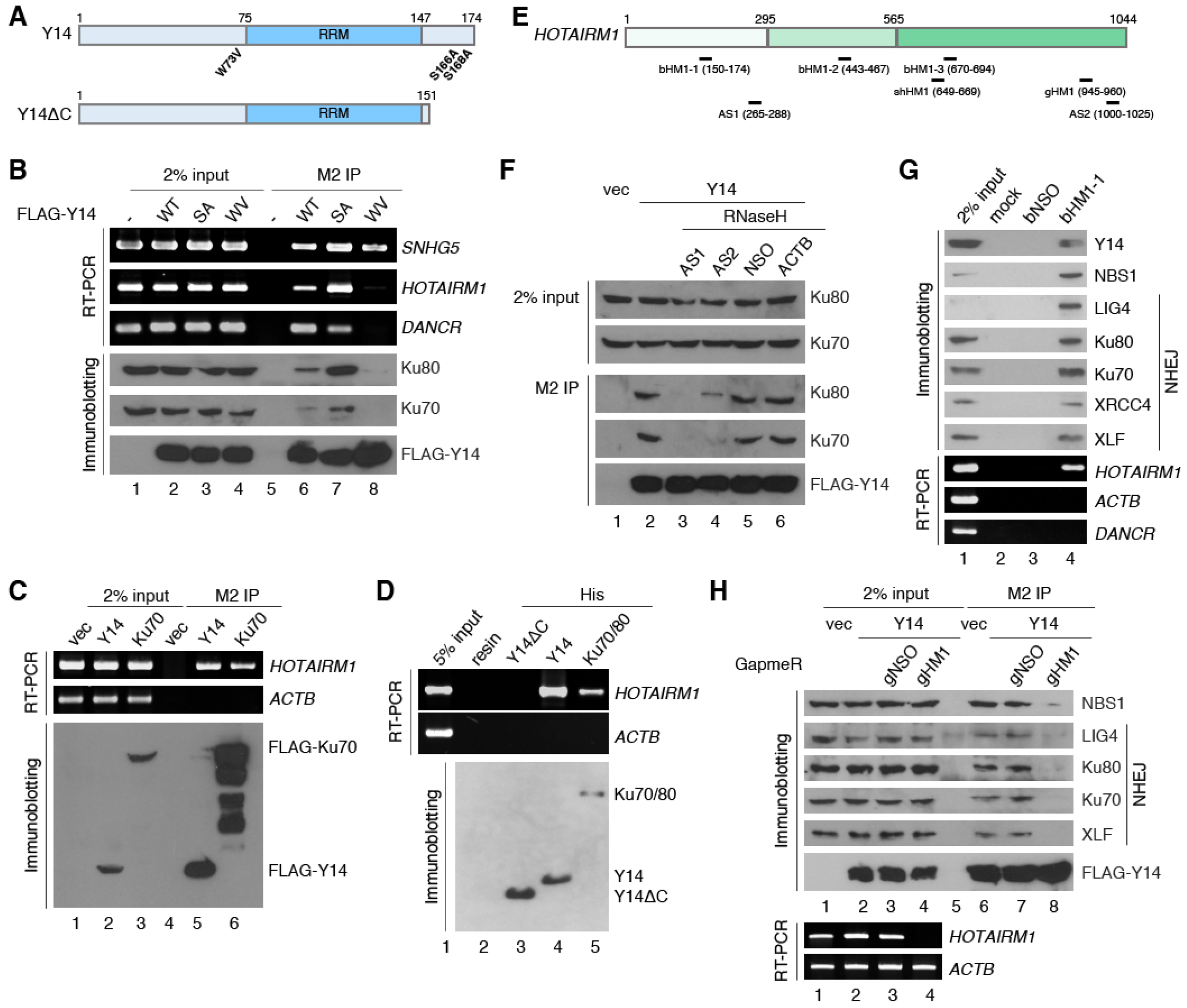
*HOTAIRM1* Is Associated with the NHEJ Complex. (A) Domains of Y14 (RRM, RNA recognition motif) and its mutants (WV and SA) and C-terminal truncated version. (B) HEK293 cells were transfected with empty (–) or FLAG-Y14-expressing vector (WT, or SA or WV mutant; WT represents wild type), followed by immunoprecipitation (IP), and RT-PCR or immunoblotting. (C) HEK293 cells were transiently transfected with empty (vec) or FLAG-Y14- or FLAG-Ku70-expressing vector, followed by immunoprecipitation along with RT-PCR or immunoblotting. (D) Recombinant His-tagged Y14, Y14ΔC or Ku70/80 was incubated with total HeLa cell RNAs, followed by pull-down using nickel resin. Bound RNAs were detected by RT-PCR. Bottom panel shows SDS-PAGE of recombinant proteins. (E) Oligonucleotides (AS, antisense; bHM, biotinylated; gHM1, GapmeR) and short hairpin RNA (shHM1) complementary to *HOTAIRM1*. (F) HEK293 cells were transfected with empty vector (vec) or FLAG-Y14 vector. Anti-FLAG immunoprecipitates were mock-treated (lane 2) or incubated with antisense oligonucleotides in the presence of RNase H (lanes 3-6). Immunoblotting was performed using antibodies against the indicated proteins. NSO and ACTB represent non-specific and β-actin-targeting antisense oligonucleotides, respectively. (G) HEK293 cell lysates were mock incubated (lane 2) or incubated with biotinylated oligonucleotides (lanes 3 and 4), followed by pull-down using strepatavidin agarose and immunoblotting or RT-PCR. bNSO: biotinylated non-specific oligonucleotide. (H) HeLa cells were transfected with empty vector (vec) or FLAG-Y14 vector or together with the indicated GapmeR. Anti-FLAG immunoprecipitates were subjected to immunoblotting and RT-PCR. gNSO: non-specific GapmeR.

### *HOTAIRM1* is Essential for the Association of Y14 with the NHEJ Complex

To investigate whether *HOTAIRM1* associates with the entire NHEJ, we performed RNA affinity selection using a biotinylated oligonucleotide complementary to *HOTAIRM1* with HEK293 cell lysates (Figures 2E and S1B, bHM1-1). bHM1-1 specifically pulled down *HOTAIRM1* as well as Y14, the DNA damage sensor NBS1, and all NHEJ factors [DNA ligase 4 (LIG4), Ku70/80, XRCC4 and XLF)] (Figure 2G and Figure S1C for the two additional biotinylated oligonucleotides tested). Next, to evaluate whether *HOTAIRM1* mediates the interaction between Y14 with the NHEJ factors *in vivo*, we transfected HEK293 cells with a *HOTAIRM1*-targeting GapmeR (Figures 2E and S1B, gHM1). Upon transfection with gHM1, *HOTAIRM1* was almost undetectable (Figure 2H, RT-PCR, lane 4). Under this condition, FLAG-Y14 no longer co-precipitated the NHEJ factors (Figure 2H, lane 8). A non-specific gNSO had no effect on the Y14-NHEJ complex (lane 7). This result emphasized that *HOTAIRM1* mediates the interaction between Y14 and the NHEJ factors.

### *HOTAIRM1* is Essential for Genome Integrity and Accumulates at DNA Damage Sites

Phosphorylation of the histone variant H2AX (γH2AX) represents a DSB marker. Depletion of *HOTAIRM1* by gHM1 increased the level of γH2AX in HeLa cells (Figure 3A, lane 4), as observed for Y14 depletion (lane 2). Next, single-cell gel electrophoresis (the comet assay) revealed that gHM1-mediated depletion of *HOTAIRM1* significantly increased the tail of cells, which reflects DNA damage (Figure 3B). All these results supported a role for *HOTAIRM1* in the maintenance of genome integrity.

**Figure 3.**
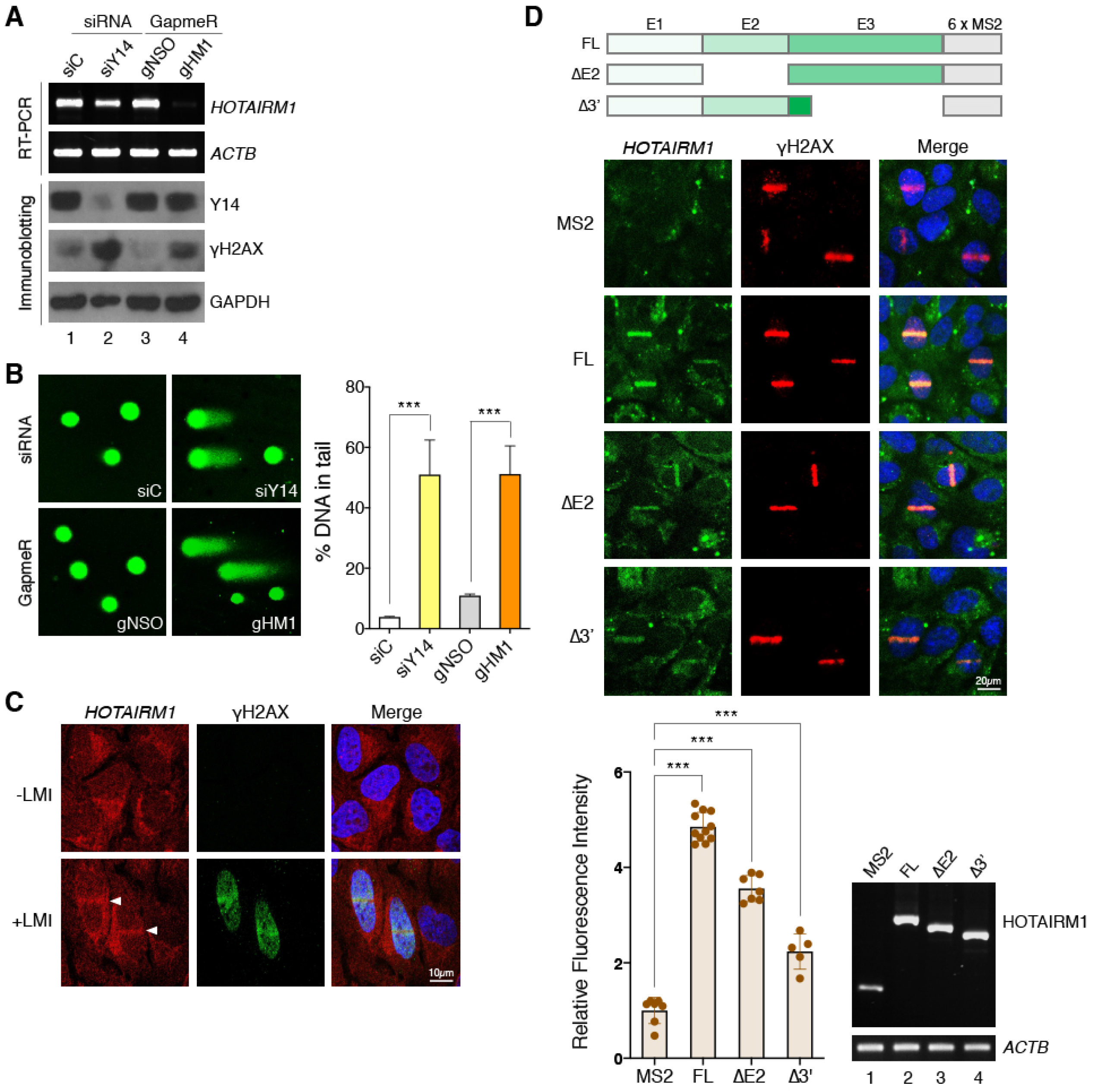
*HOTAIRM1* Accumulates at DNA Damage Sites and Contributes to Genome Integrity. (A) HeLa cells were transfected with indicated siRNA or GapmeR. RT-PCR and immunoblotting were performed 48 h post-transfection. (B) HeLa cells were transfected as in panel A, followed by the comet assay. Bar graph shows the percentage of DNA in the comet tail (mean±SD); \*\*\**p* < 0.001. (C) U2OS cells were mock-irradiated (–LMI) or laser microirradiated (405 nm; +LMI) and then immediately fixed for *in situ* hybridization using bHM1-3 as the probe and subsequently subjected to immunofluorescence microscopy using anti-γH2AX. (D) *HOTAIRM1*-MS2 chimeric RNA and truncations. E indicates exon. U2OS cells were transfected with the control vector (MS2 only) or a *HOTAIRM1*-MS2-expressing vector and the GFP-MCP-expressing vector (GFP signals represent *HOTAIRM1*), followed by indirect immunofluorescence microscopy using anti-γH2AX. Dot graph shows the relative GFP/γH2AX intensities (MS2 was set to 1); \*\*\**p* < 0.001.

Next, we examined whether endogenous *HOTAIRM1* is located at DNA damage sites. Use of bHM1-3 as a probe for *in situ* hybridization revealed that *HOTAIRM1* distributed primarily in the cytoplasm but also the nucleus of U2OS cells, as previously reported (Rea et al., 2020) (Figure 3C, without laser microirradiation, -LMI). LMI induced *HOTAIRM1* accumulation at DNA damage tracks, as indicated by γH2AX (Figure 3C, +LMI). To confirm the recruitment of *HOTAIRM1* to DNA damage sites, we tagged *HOTAIRM1* with six repeats of the RNA stem-loop of bacteriophage MS2, which can be recognized by the MS2 capsid protein (MCP). U2OS cells were co-transfected with a vector expressing MS2 only or *HOTAIRM1*-MS2 and GFP-MCP. Indirect immunofluorescence microscopy revealed that *HOTAIRM1*-MS2 but not M2S localized to LMI-induced damage tracks [Figure 3D, FL (full-length) and MS2]. During the course of this study, we found that *HOTAIRM1* existed in two isoforms, *i*.*e*. full-length and exon 2-skipped isoform (ΔE2). Their expression ratio differed between cell lines; the latter was dominant in U2OS cells (Figure S2A). Nevertheless, both could associate with Y14 (Figure S2B). MS2-tagged ΔE2 accumulated at DNA lesions after LMI, albeit with less efficiency than the full-length isoform (Figure 3D, bar graph). Truncation of 431 nucleotides from the 3’ end (Δ3’) more severely affected *HOTAIRM1* localization. Together, all the results indicated that *HOTAIRM1* is recruited to DNA damage sites.

### *HOTAIRM1* Modulates the Recruitment and Retention of DNA Repair Factors at DNA Damage Sites

Next, we took advantage of live-cell imaging of U2OS cells that stably expressed a GFP fusion with MDC1 or Ku70/80 (Chang et al., 2015) to evaluate the effect of *HOTAIRM1* in localization of repair factors to laser-induced DNA damage sites. These U2OS cells were transiently transfected with a vector expressing a *HOTAIRM1*-targeting short hairpin RNA (shHM1) (Figures 2E and S1B) and RFP, which served as a marker of transfected cells. shHM1 downregulated *HOTAIRM1* by up to 70% (Figure 4A, RT-PCR). LMI caused a gradual accumulation of GFP-tagged DNA damage repair factors (MDC1, Ku70 and Ku80) at DNA damage sites in mock-transfected cells, and this accumulation was considerably reduced in shHM1-expressing (RFP-positive) cells (Figure 4A, fluorescence live-cell imaging and line graphs). This result indicated that *HOTAIRM1* is required for the recruitment of DNA repair factors. Depletion of Y14 leads to Ku accumulation on chromatin after DNA damage(Chuang et al., 2019). Therefore, we examined whether *HOTAIRM1* depletion affects the IR-induced formation of Ku70 foci. Use of an RNase A-based extraction method revealed that IR transiently increased the signal of Ku70 foci in U2OS cells (Figure 4B, gNSO for 5 and 30 min), as previously reported (Brown et al., 2015). The Ku70 signal markedly increased from 5 to 60 min after IR treatment and returned to baseline at 2 h in HOTAIRM1-depleted cells (Figure 4B, gHM1). The kinetics of γH2AX foci formation did not significantly differ between control and *HOTAIRM1*-depleted cells (Figures 4B and S3, γH2AX). This result was similar to that observed with inhibition of Ku ubiquitination during DNA damage repair (Brown et al., 2015). Therefore, *HOTAIRM1* depletion may cause Ku retention or impair Ku removal from DNA repair sites. Together, these results suggested that *HOTAIRM1* promotes efficient loading of the repair factors to DSBs and is also required for their dissociation from DSBs.

**Figure 4.**
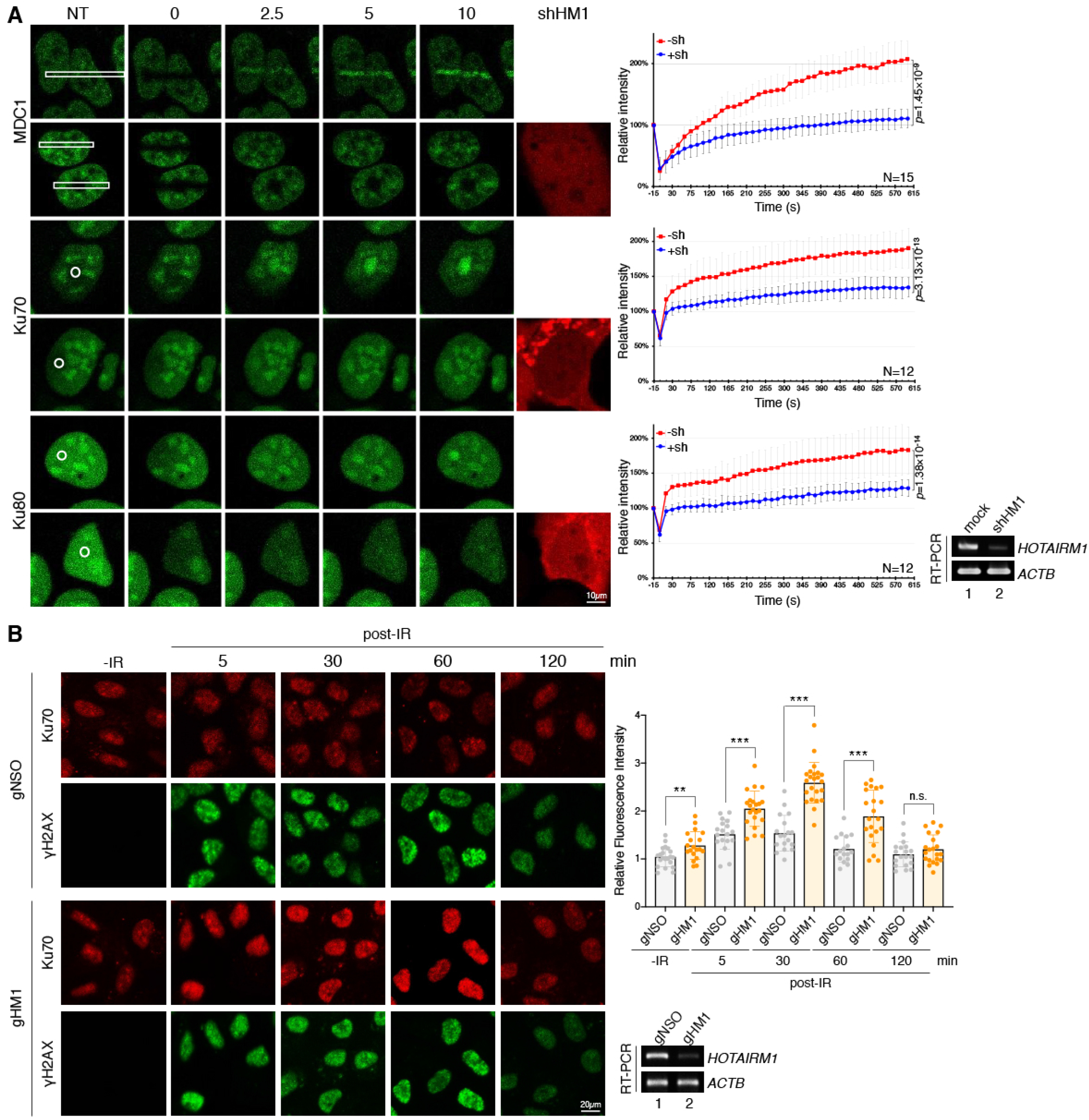
*HOTAIRM1* Is Required for the Recruitment of DNA Repair Factors to DSB sites and Regulates Ku foci dynamics. (A) U2OS cells that stably expressed the GFP fusion with MDC1, Ku70, or Ku80 were mock-transfected or transfected with the shHORAITM1 (shHM1)-mCherry-expressing vector. Cells were subjected to laser microirradiation followed by live-cell imaging using confocal microscopy. Representative confocal images show accumulation of GFP fusion proteins at sites (white-lined rectangles or circles) of laser microirradiation at the indicated time points. NT (non-treated) indicates samples before microirradiation. mCherry represents sh*HOTAIRM1*-expressing cells. Graphs to the right show fluorescence intensities of GFP-fusion proteins at the irradiated region. Intensity was quantified periodically up to 10 min, normalized, and is presented as the mean and standard deviation (*p*-value as indicated) for 12 or 15 cells in each experiment. The RT-PCR data indicate the efficiency of *HOTAIRM1* knockdown. (B) U2OS cells were transfected with gNSO or gHM1. Cells were not irradiated (–IR) or exposed to 10 Gy of IR and harvested at the indicated time points post-IR. Indirect immunofluorescence microscopy was performed using anti-γH2AX and anti-Ku70. Dot graph shows relative fluorescence intensity of Ku70; for each sample, 16-22 cells were measured (–IR was set to 1); \*\*\**p* < 0.001.

### *HOTAIRM1* is Required for DSB Repair

Next, we adopted two assay systems to evaluate whether *HOTAIRM1* is essential for DSB repair. First, using CRISPR/Cas9-mediated DNA cleavage at a specific gene, namely hypoxanthine guanine phosphoribosyltransferase (HPRT), we analyzed DNA repair in the presence of a blunt-ended double-stranded oligonucleotide Ins (Du et al., 2018) (Figure 5A). Insertion of Ins into the cleavage site, indicating DNA repair (Du et al., 2018; Tsai et al., 2015), was evaluated by PCR using primers complementary to Ins (primer I) and the region downstream of the cleavage site (primer R). The PCR product I-R was detectable upon transfection of cells with both the Cas9/sgHPRT vector and Ins (Figure 5B, upper, lane 3) but not detected by transfection with either (lanes 2 and 9). Quantitative PCR (qPCR) revealed that knockdown of Y14 or LIG4 using a small interfering RNA (siRNA) reduced the efficiency of DNA repair by ∼50% and 70%, respectively, and *HOTAIRM1* depletion impaired DNA repair by ∼40% (Figure 5B, bar graph). To confirm this observation, we additionally used HeLa NHEJ reporter cells for analysis (Allen et al., 2017). The chromosomally integrated GFP reporter gene was disrupted by an intron, within which an inserted exon (Ad2) was flanked by two I-SceI sites (Figure 5C). Transfection of cells with the I-SceI expression vector (pSCE) induced cleavage, which mimicked a DSB. Upon DNA repair, the GFP reporter gene was expressed, producing the transcript, within which Ad2 had been removed by splicing. In general, ∼5% GFP-positive cells were observed in pSCE-transfected cells. Knockdown of Y14 or *HOTAIRM1* reduced the number of GFP-positive cells by 36% and 53%, respectively (Figure 5D, bar graph). This result further supported the role of *HOTAIRM1* in DSB repair likely via the NHEJ pathway.

**Figure 5.**
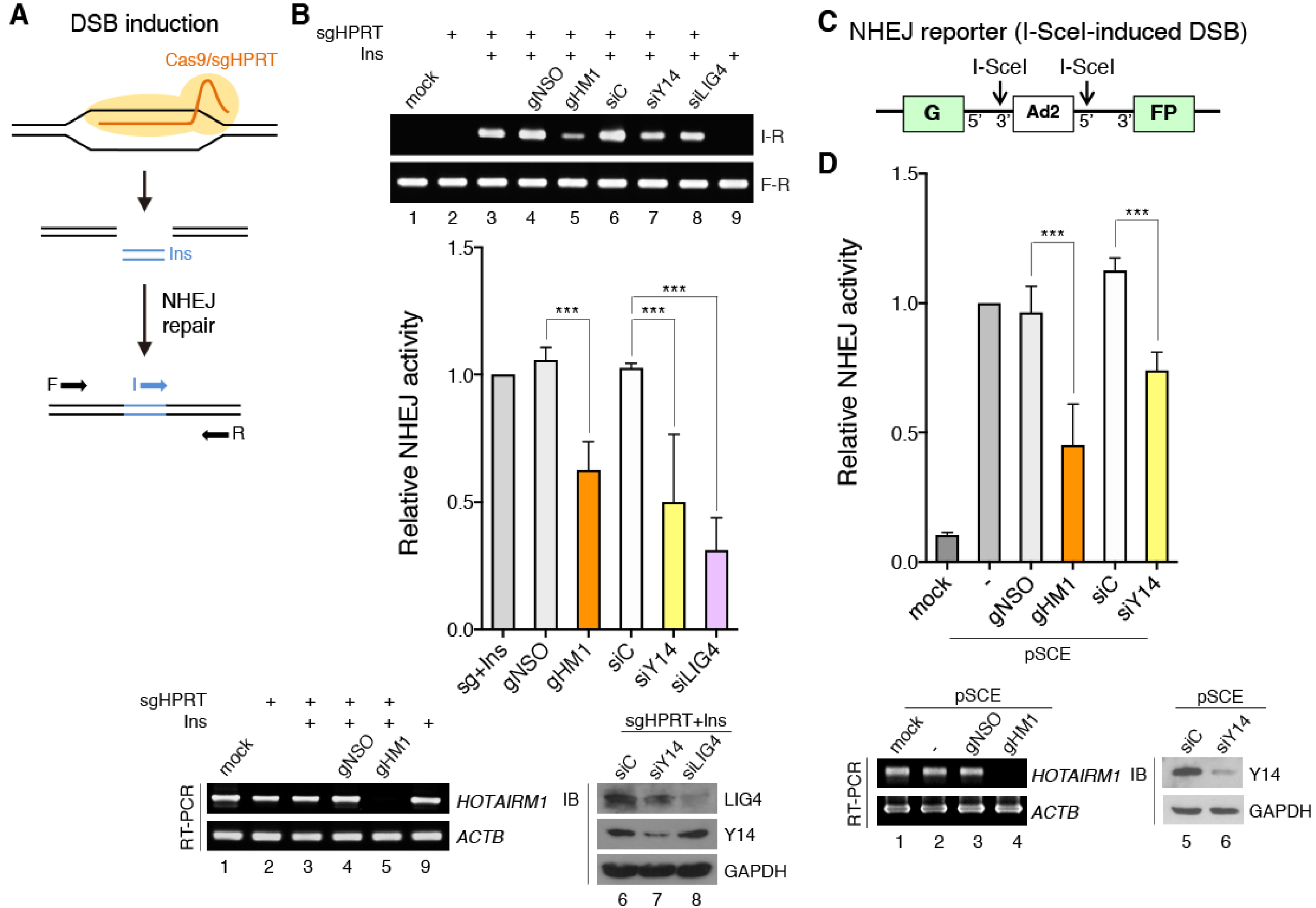
*HOTAIRM1* Is Required for DSB Repair. (A) Experimental design for quantitative measurement of NHEJ-mediated repair of DSBs. Transfection of HeLa cells with the Cas9/sgRNA (sgHPRT) vector induces DSBs. Incorporation of the double-stranded oligonucleotide ‘Ins’ into the DSB sites was evaluated by PCR and qPCR represent DNA repair efficiency. (B) HeLa cells were mock-transfected (lane 1) or transfected for 48 h with one or more of the following reagents: the Cas9/sgHPRT plasmid, Ins, and GapmeR (lanes 4 and 5) or siRNA (lanes 6–8) as indicated. Genomic DNA was recovered for PCR using the primer set I/R or F/R. Bar graph shows qPCR of lanes 3-8 (sg+Ins was set to 1). Immunoblotting and RT-PCR indicate the knockdown efficiency; lanes are numbered to correspond with the PCR analysis (upper panel). (C) HeLa NHEJ reporter cells in which the chromosomally integrated GFP gene is disrupted by an intron and flanked by I-SceI sites. I-SceI-induced cleavage mimics a DSB. GFP expression was restored after the cleavage repaired through NHEJ. (D) HeLa NHEJ reporter cells were mock-transfected or transfected with pSCE (the I-SceI expression vector) and GapmeR or siRNA as indicated. The number of GFP-positive cells was counted 48 h post-transfection. Bar graph represents relative repair efficiency; samples without GapmeR/siRNA transfection were set to 1. Immunoblotting and RT-PCR show knockdown efficiency. For the bar graphs in panels B and D, average and standard deviation were obtained from 3-5 independent experiments; \*\*\**p* < 0.001.

### *HOTAIRM1* Associates with DNA Damage Repair Factors and RNA Processing Factors

To gain further insight into *HOTAIRM1*-mediated DNA repair, we affinity-selected *HOTAIRM1* in the lysates of UV-crosslinked HEK293 cells using three bHM1 oligonucleotides (Figure 6A). Co-purified proteins were gel-fractionated and analyzed by liquid chromatography-tandem mass spectrometry. Among the 3990 proteins that were identified (Table S3), 688 proteins with ≥5 protein member hits in the MASCOT search results were subjected to pathway enrichment analysis with Kyoto Encyclopedia of Genes and Genomes (KEGG) using DAVID software (david.ncifcrf.gov) (Table S5). The results indicated that *HOTAIRM1*-associated proteins are primarily involved in RNA processing and DNA damage repair (Figure 6B). Spliceosomal and ribosomal factors were at the top of the ranked list (respective *p*-values, 6.35E-22 and 1.44E-20). In addition, a set of mRNA processing and surveillance factors were identified, some of which have been implicated in DNA damage repair, such as the RNA exosome component Exosc10 (Rrp6) (Domingo-Prim et al., 2019; Marin-Vicente et al., 2015). The most highly represented DNA damage repair pathways were NHEJ and HR (respective *p*-values, 0.022 and 0.049). Moreover, we arbitrarily selected 166 proteins that form functional complexes involved in transcription, RNA processing, DNA replication or repair, or genome stability for analysis of protein-protein interaction networks using STRING (string-db.org) (Table S5). The results underscored *HOTAIRM1-*mediated connection between transcription, RNA processing, and various DNA repair pathways (Figure 6C). Moreover, identification of PARP1, the MRN complex, and four NHEJ factors (Ku80, Ku70, DNA-PK and LIG4) supported the role of *HOTAIRM1* in the NHEJ pathway. Nevertheless, we cannot exclude the possibility that *HOTAIRM1* participates in other DNA repair pathways (Figure 6C and Table S6). Notably, we identified a set of NMD factors, namely Upf1, Upf3B, SMG5, SMG6, SMG7, SMG8 and SMG9. Although NMD factors function in cytoplasmic mRNA surveillance, some of them, such as Upf1 and SMG6, have been implicated in genome/telomere integrity (Azzalin and Lingner, 2006; Reichenbach et al., 2003; Snow et al., 2003). Moreover, a recent report has revealed a role for Upf1 in HR-mediated DSB repair (Ngo et al., 2021). Using affinity selection, we confirmed the association of these two factors with *HOTAIRM1*, whereas the unidentified factor SMG1 was not detected (Figure 6D). Moreover, the interaction between *HOTAIRM1* and any of Y14, DNA repair factors, or Upf1/SMG6 was independent of DNA damage (Figure S4A). Y14, however, interacted directly with Upf1, suggesting that Y14 recruits mRNA surveillance factors to *HOTAIRM1* (Figure S4B).

**Figure 6.**
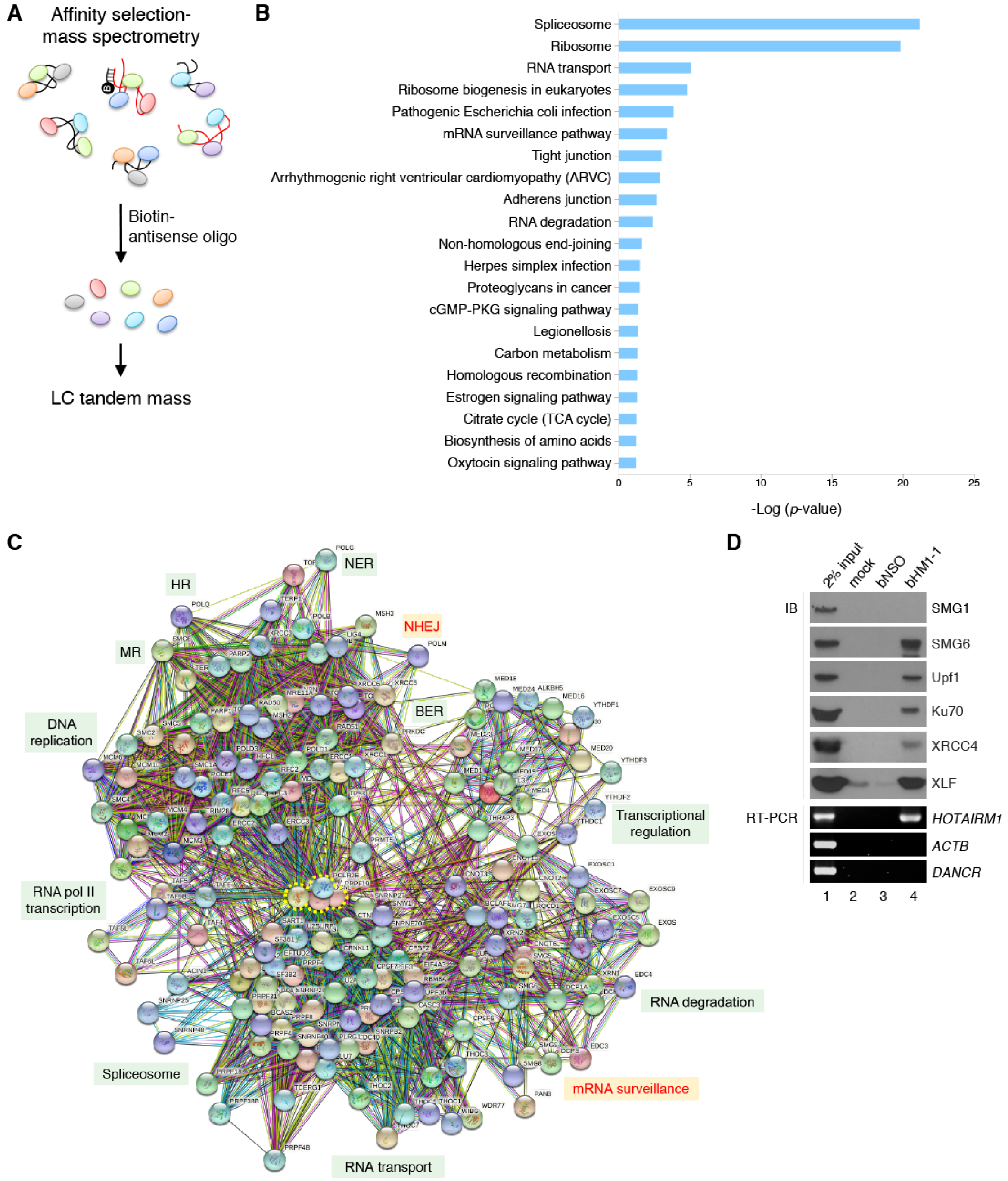
*HOTAIRM1* Is Associated with the mRNA Surveillance Factors. (A) Procedure for identification of *HOTAIRM1*-interacting proteins. HEK293 cell lysates were incubated with three bHM1 oligonucleotides (see STAR Methods) followed by affinity selection using streptavidin. Selected proteins were analyzed by mass spectrometry. (B) KEGG pathway enrichment analysis of the 688 *HOTAIRM1* partners that were identified. Bar graph shows the top enriched Gene Ontology terms (*p*-value < 0.05) from the KEGG Pathway Database. (C) STRING analysis of the 166 *HOTAIRM1*-interacting proteins (Table S6) that function in gene expression, DNA replication, or DNA repair. Diagram shows protein-protein interaction (PPI) networks for these proteins (PPI enrichment *p*-value < 1.0e-16). NHEJ and mRNA surveillance are highlighted in red. RNA polymerase II (RNA pol II) subunits and the PRP19-CDCL5 complex are enclosed by a dotted yellow line and represent hub proteins. BER, base excision repair; MR, mismatch repair; NER, nucleotide excision repair. (D) Affinity selection of *HOTAIRM1* was as in panel A. RT-PCR and immunoblotting were performed to detect *HOTAIRM1* and its interacting partners.

### The mRNA Surveillance Factors Upf1 and SMG6 Participate in DSB Repair

Next, we investigated whether Upf1 and SMG6 participate in DNA damage repair. Both Upf1 and SMG6 were found to reside predominantly in the cytoplasm of U2OS cells (Figure 7A, –IR). To our surprise, IR induced drastic nuclear translocation of SMG6, although Upf1 was still largely retained in the cytoplasm (Figure 7A, +IR). Nevertheless, both Upf1 and SMG6 localized to laser-induced DNA damage tracks (Figure 7B; transfection with gNSO), suggesting their potential involvement in DNA damage repair. Noteworthily, depletion of *HOTAIRM1* abolished the laser-induced localization of Upf1 and SMG6 at DNA damage sites (Figure 7B; gHM1), indicating that *HOTAIRM1* escorts mRNA surveillance factors to DNA lesions. Next, using Cas9/sgHPRT-induced DNA cleavage, we assessed whether these two factors have a role in DNA repair. Depletion of Upf1 or SMG6 by siRNA reduced DNA repair efficiency by ∼60% and 35%, respectively (Figure 7C), indicating that both are required for efficient DSB repair.

**Figure 7.**
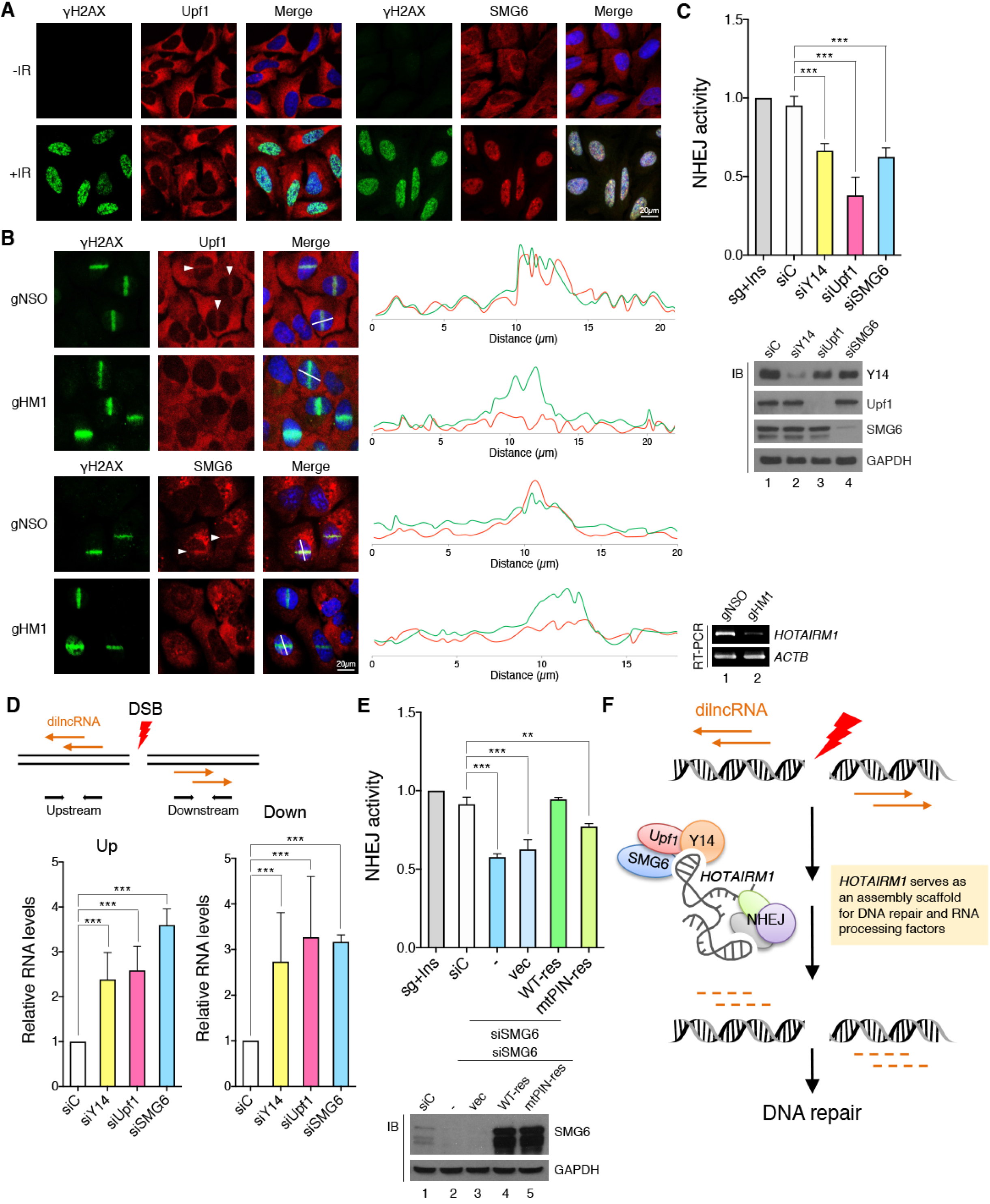
mRNA Surveillance Factors Are Required for DSB Repair. (A) U2OS cells were mock mock-irradiated (–IR) or irradiated with 10 Gy (+IR) followed by immunofluorescence microscopy using antibodies against the indicated proteins. (B) U2OS cells were transfected with gNSO or gHM1. LMI was performed, followed by immunofluorescence microscopy using antibodies against the indicated proteins. Arrowheads indicate laser-irradiated cells. Fluorescence intensities along a white line across a nucleus were measured in arbitrary units. Line-scan profiles of fluorescence intensity are shown to the right. RT-PCR shows *HOTAIRM1* knockdown efficiency. (C) The DSB repair assay was performed as in Figure 5B. HeLa cells were transfected with the Cas9/sgHPRT vector, Ins, and siRNA as indicated. Genomic DNA was collected at 48 h post-transfection and subjected to qPCR. Bar graph is shown as in Figure 5B; \*\*\**p* < 0.001. (D) Schematic drawing of Cas9/sgHPRT-generated DSBs and production of dilncRNAs in the DSB-flanking regions. HeLa cells were transfected similarly as in panel C. RT-qPCR was performed using the primers indicated in the diagram. Bar graphs show relative levels of dilncRNAs (mean±SD); \*\*\**p* < 0.001. (E) HeLa cells were transfected with the Cas9/sgHPRT vector, Ins, siC, or siSMG6 together with the empty vector (vec) or the siRNA-resistant wild-type or mutant SMG6 expression vector (WT-res or mtPIN-res). qPCR and immunoblotting were performed as in panel C; \*\*\**p* < 0.01. (F) Model showing that *HOTAIRM1* associates with DNA repair factors and RNA processing factors. Upon DNA damage, *HOTAIRM1* accumulates at DSBs and is essential for efficient recruitment of Ku70/80 and subsequent DSB repair. *HOTAIRM1* also recruits mRNA surveillance factors to DSBs, which regulates dilncRNA turnover. dilncRNA accumulation may impede DNA repair.

DSB-induced dilncRNAs are bidirectionally synthesized, and their turnover is in part regulated by Exosc10 (Domingo-Prim et al., 2019; Marin-Vicente et al., 2015). The mRNA surveillance machinery has a role in degrading nonsense mRNAs in the cytoplasm (Kurosaki et al., 2019). We examined whether it also regulates the abundance of dilncRNAs at DNA damage sites. RT-qPCR using primers complementary to sites upstream or downstream of the Cas9/sgHPRT-induced cleavages revealed that depletion of Y14 or either of Upf1 or SMG6 increased the level of dilncRNAs by at least 2-3 folds (Figure 7D). Therefore, the mRNA surveillance machinery may participate in the degradation of the transcripts generated from DSBs. Because SMG6 has endonucleolytic activity, we further analyzed whether this activity is essential for DNA repair. Overexpression of siRNA-resistant SMG6 almost fully restored DNA repair in SMG6-depleted cells (Figure 7E, WT-res). The PilT N-terminus (PIN) domain in the C-terminal region of SMG6 is important for its catalytic activity in NMD (Glavan et al., 2006). Overexpression of a SMG6-PIN mutant (D1251A) (Boehm et al., 2014; Okada-Katsuhata et al., 2012) only partially restored DNA repair (Figure 7E, mtPIN-res), suggesting that SMG6 participates in DSB repair at least in part by degrading dilncRNAs at DSBs.

Together, these results revealed that the mRNA surveillance factors recruited to DSBs by *HOTAIRM1* participate in dilncRNA turnover, which influences the efficiency of DSB repair.

## DISCUSSION

### The Role of *HOTAIRM1* in DSB Repair

Our present study reveals that *HOTAIRM1* contributes to DSB repair via its interaction with Y14, mRNA surveillance factors, and NHEJ factors (Figure 7F). DSB repair involves different types of RNA, including dilncRNA and their processed small RNAs and pre-existing lncRNAs (Domingo-Prim et al., 2020; Hawley et al., 2017; Wu and Wang, 2017). Among Y14-associated lncRNAs, *HOTAIRM1* fulfilled criteria making it a likely participant in Y14-mediated DNA damage repair (Figure 1). At present, we cannot exclude any other candidates involved. For example, two *SNHG* lncRNAs were recently reported to contribute to DSB repair (Haemmig et al., 2020; Han et al., 2020). Nevertheless, the association of both Y14 and eIF4A3 with the *SNHG* RNAs we examined suggests that they participate in *SNHG* RNA processing and/or surveillance. In this study, we provide several lines of evidence for the role of *HOTAIRM1* in DSB repair. First, *HOTAIRM1* interacted with Y14 and NHEJ factors (Figure 2). LMI caused *HOTAIRM1* to localize to DNA damage sites (Figure 3). *HOTAIRM1* was essential for efficient recruitment of NHEJ factors and also regulated Ku70 foci dynamics (Figure 4). Finally, *HOTAIRM1* recruited Upf1/SMG6 to DSB sites, which regulate dilncRNA turnover (Figure 7).

The association of *HOTAIRM1* with the NHEJ factors is reminiscent of several lncRNAs that have been implicated in NHEJ (Haemmig et al., 2020; Han et al., 2020; Unfried et al., 2021; Zhang et al., 2016). *LINP1* was the first lncRNA identified, and it acts as a modular scaffold linking Ku and DNA-PK (Zhang et al., 2016). Another study revealed that *LINP1* self-assembles into phase-separated condensates that sequester Ku, and this stabilizes the initial interaction between DNA ends during synapsis and repair (Thapar et al., 2020). Similar to *LINP1, NIHCOLE* forms clusters with Ku and promotes ligation of DSBs via recruitment of several NHEJ factors (Unfried et al., 2021). Therefore, potential *NIHCOLE* phase separation may favor repair kinetics. We found that *HOTAIRM1* could bind recombinant Y14 and Ku directly *in vitro* (Figure 2). Therefore, it would be interesting to determine the nature of the molecular interactions between *HOTAIRM1* and Y14/Ku. Finally, STRING analysis of the *HOTAIRM1* interactome revealed a number of factors involved in mRNA biogenesis. In particular, RNA polymerase II and the PRP19-CDC5L complex act as a hub. PRP19 not only functions in precursor mRNA splicing but also plays a critical role in multiple DDR signaling networks (Mahajan, 2016; Marechal et al., 2014). Notably, the lncRNA *NORAD* complex also contains PRP19 and contributes to genomic stability (Munschauer et al., 2018). Therefore, whether certain lncRNAs, such as *NORAD* and *HOTAIRM1*, share some common factors to coordinate RNA processing and DNA repair warrants further investigation.

Besides pre-existing lncRNAs, DSB-induced dilncRNAs also participate in DNA repair by forming RNA-RNA or RNA-DNA hybrids at DSB sites (Chakraborty et al., 2016; D’Alessandro et al., 2018; Michelini et al., 2017). dilncRNAs form RNA-DNA hybrids with the resected DNA ends during S/G2 phase of the cell cycle. After degradation of RNA by RNase H2, Rad51 is loaded onto the resultant single-stranded DNA for HR-based repair (D’Alessandro et al., 2018). On the other hand, dilncRNAs can be processed into small RNAs in repetitive regions or ribosomal DNA loci via DICER-dependent or -independent pathways (Bonath et al., 2018; Michelini et al., 2017). RNA hybrids formed by dilncRNAs and those small RNAs can promote the formation of 53BP1-containing DDR foci (Michelini et al., 2017). Recent evidence indicates that phase-separated condensates of DDR factors provide transient repair compartments for efficient DNA repair (Pessina et al., 2019). RNA itself has the potential to drive phase separation of DDR factors. For example, inhibition of RNA polymerase II-mediated transcription or disruption of RNA condensation prevents the formation of 53BP1 foci, suggesting a role for dilncRNAs in promoting molecular crowding of DNA repair factors (Pessina et al., 2019). Therefore, *HOTAIRM1* may not only deliver factors to DNA damage sites for RNA processing and DNA repair but also self-assemble into phase-separated condensates, as does *LINP*, to facilitate the repair reaction; the latter remains to be clarified by future investigation (Thapar et al., 2020).

### The Role of mRNA Surveillance Factors in DSB Repair

Using RNA affinity selection, we identified Upf1 and SMG6 as *HOTAIRM1*-interacting partners (Figure 6). Upf1 and SMG6 primarily function in NMD in the cytoplasm. Upon association with a stalled ribosome on nonsense mRNAs, Upf1 is phosphorylated by the phosphatidylinositol 3-kinase-related kinase (PIKK) SMG1 and subsequently recruits SMG6 and SMG5/7 to degrade RNA (Chakrabarti et al., 2014; Okada-Katsuhata et al., 2012). SMG1 was not detected in the *HOTAIRM1* ribonucleoprotein complex (Figure 6). Therefore, the possibility that the DDR-related PIKK kinases such ATM and ATR phosphorylate Upf1 at DNA damage sites should be investigated. Several early studies have revealed the potential role of Upf1 and SMG6 in maintaining genome/telomere integrity. Upf1 associates with chromatin during S phase and after DNA damage, whereas SMG6 is a telomerase cofactor (Azzalin and Lingner, 2006; Reichenbach et al., 2003; Snow et al., 2003). Notably, a recent report indicated that Upf1 can promote DNA resection and repair at subtelomeric DSBs by driving R-loop formation (Ngo et al., 2021). Our result that both Upf1 and SMG6 located to laser-induced DNA damage tracks and that SMG6 shifted from the cytoplasm into the nucleus upon IR treatment emphasizes their general role in DNA damage repair (Figure 7).

Several ribonucleases have been implicated in DNA damage repair. An early study indicated that the yeast exonuclease XRN1 promotes end resection probably by degrading RNAs in the vicinity of DSBs (M8). XRN2 plays a role in R-loop resolution and genomic stability (M9). More recent studies indicated that DICER and RNase H2, respectively, participate in dilncRNA processing into small RNAs or dilncRNA degradation during S/G2 phase (M4, D5). Exosc10 degrades dilncRNAs released from DNA:RNA hybrids by the helicase senataxin, leading to replication protein A loading onto resected DNA ends (Cohen et al., 2018; Domingo-Prim et al., 2019). Therefore, the function of the RNA helicase Upf1 and the endonuclease SMG6 in dilncRNA turnover may be analogous to that of senataxin and Exosc10. Moreover, depletion of Upf1 or SMG6 increased the level of dilncRNAs (Figure 7). We postulate that dilncRNA accumulation hampers Ku dissociation from DSBs (Figure 4), which may be similar to the scenario in which the prevention of ubiquitination or phosphorylation of Ku70/80 abolishes the dissociation of Ku70/80 from DSBs and hence inhibits break repair (Ishida et al., 2017; Lee et al., 2016a). Therefore, dilncRNA clearance is likely a necessary step for DSB repair. Whether different ribonucleases function in different repair pathways needs future investigation.

## ACKNOWLEDGEMENTS

We are grateful for Niels Gehring (University of Cologne, Germany) and Shigeo Ohno and Akio Yamashita (Yokohama City University, Japan) for plasmids. We thank the Proteomics Core Facility and Light Microscopy Core Facility of the Institute of Biomedical Sciences, and NGS High Throughput Genomics Core Facility, Academia Sinica. We appreciate Shih-Han Ko, Chun-Hao Su and Wei-Chi Ke for constructing vectors and technical assistance. This work was supported by Academia Sinica Investigator Award (AS-IA-107-L04) and Grant 109-2811-B-001-529 of the Ministry of Science and Technology, Taiwan, to W.Y.T.

## AUTHOR CONTRIBUTIONS

T.W.C. conducted the majority of the experiments and wrote STAR Methods. P.Y.W. performed laser microirradiation and time laps imaging (Figure 4A). Y.M.C. carried out RNA seq analysis (Figure 1B-E, Tables S1-S2). W.Y.T. conceived the project, supervised the experiments, performed bioinformatics analysis (Figure 6B and 6C, Tables S3-S6) and wrote the manuscript.

## SUPPLEMENTAL FIGURE LEGENDS

**Figure S1. Characterization of Y14 and *HOTAIRM1*. Related to Figure 2**.

(A) HEK293 cells were transiently transfected with the vector expressing FLAG-Y14 (lanes 1 and 2) or FLAGY14-SA (lanes 3 and 4). Cell fractionation was performed as previously described (Chuang et al., 2019). Immunoblotting was performed using antibodies against indicated proteins. T, total; C, chromatin fraction.

(B) Secondary structure of *HOTAIRM1* was predicted by RNAfold Webserver (rna.tbi.univie.ac.at). All types of antisense oligonucleotides used in this study are indicated as in Figure 2E.

(C) Affinity selection of *HOTAIRM1* was performed as in Figure 2G using three different bHM1 oligonucleotides. Immunoblotting and RT-PCR was performed as in Figure 2G.

**Figure S2. Expression of Two Isoforms of *HOTAIRM1*. Related to Figure 3D**.

(A) RT-PCR was performed in cell lines as indicated using primers complementary to exons 1 and 3.

(B) Immunoprecipitation of FLAG-Y14 from HEK293 cell lysates, RT-PCR and immunoblotting were performed as in Figure 1F.

**Figure S3. Depletion of *HOTAIRM1* does not Alter** γ**H2AX foci formation. Related to Figure 4B**.

U2OS cells were transfected and irradiated as in Figure 4B. Bar graph shows the relative intensity of γH2AX foci (gNSO was set to 1).

**Figure S4. Characterization of the Interaction Between *HOTAIRM1* and DNA Repair and RNA Processing Factors. Related to Figure 6D**.

(A) HEK293 cells were mock-irradiated (–IR) or irradiated (10 Gy; +IR). Cell lysates were incubated with bNSO and bHM1, followed by affinity selection using streptavidin. Immunoblotting and RT-PCR were performed using specific antibodies or primers as indicated.

(B) HEK293 cells were transfected with the empty or FLAG-Y14 vector. FLAG-Y14-containing cell lysates were mock-treated or treated with RNase A. Cell lysates (vec or FLAG-Y14 without or with RNase A treatment) were subjected to immunoprecipitation using anti-FLAG. Immunoblotting was performed using antibodies against indicated proteins.

## STAR Methods

### Cell Culture and Transfection

Cell culture and transient transfection of HeLa, HEK293 and U2OS cells were performed as previously described (Chuang et al., 2019). All siRNAs, antisense oligonucleotides, biotinylated antisense oligonucleotides, antisense GapmerRs are listed in Table S8. U2OS cells that stably express GFP fusion DDR factors (Ku80, Ku70 or MDC1) were used previously (Chang et al., 2015; Chuang et al., 2019).

### Plasmids

The expression vectors encoding FLAG-tagged Y14 (wild-type, SA or WV mutant), eIF4A3 and Ku70 were described previously (Chuang et al., 2019). The pLKO.1-shHOTAIRM1-mCherry vector was constructed as previously described for pLKO.1-shY14-mCherry (Chuang et al., 2019); the shHOTAIRM1 sequence was: 5’-AAATGTGGGTGTTTGAAACAACTCGAGTTGTTTCAAACACCCACATTT. The pSCE expression vector was provided by the HeLa cell-based NHEJ screening kit (TopoGEN). For CRISPR/Cas9-mediated DNA cleavage, the pAll-Cas9.Ppuro vector expressing an HPRT gene-targeting sgRNA (5’-GCAAAGGGTGTTTATTCCTCA-3’) was constructed according to Du et al. (Du et al., 2018) by National RNAi Core Facility, Academia Sinica, Taiwan. The *HOTAIRM1* cDNA was chemically synthesized (TOOLS, Taiwan), and inserted into pcDNA 3.1. Three stop codons and six MS2-binding sites that were PCR-amplified from the β6MS2 reporter (Hsu Ia et al., 2005) were inserted into the 5’ and 3’ ends, respectively, resulting in pcDNA-HOTAIRM1-6×MS2. The truncated versions of *HOTAIRM1* were generated by using a PCR-based strategy. The pcDNA-MCP-GFP expression vector was described previously (Hsu Ia et al., 2005). The SMG6-expressing vector was a kind gift of Shigeo Ohno and Niels Gehring (Boehm et al., 2014; Okada-Katsuhata et al., 2012). The SMG6-PIN mutant was generated by PCR-based mutagenesis. The sequence of all the resulting constructs was confirmed.

### Chromatin Fractionation and Immunoprecipitation

HEK293 cells were transiently transfected with an empty vector or FLAG-tagged Y14. Forty-eight hours post-transfection, cells were washed with phosphate buffered saline (PBS) and then incubated at 4°C for 3 min in the cytoskeleton (CSK) buffer containing 10 mM PIPES (pH 7.0), 100 mM NaCl, 300 mM sucrose, 3 mM MgCl_2_, and 0.7% Triton X-100. The chromatin-enriched pellet was removed by centrifugation, washed with CSK buffer twice, resuspended in CSK buffer containing 500 mM NaCl, and finally incubated at 4°C for 15 min. After centrifugation at 12,000 × *g* for 15 min, the supernatant was collected as the chromatin-associated fraction and used for immunoprecipitation. For immunoprecipitation, the lysates were incubated with anti-FLAG M2 affinity gel (Sigma) at 4°C for 2 hours. Beads were then washed with NET-2 buffer (150 mM NaCl, 50mM Tris-HCl and 0.05% NP-40), followed by RNA extraction using TRIzol reagent (Thermo Fisher Scientific) or protein extraction using SDS-PAGE sample buffer (Chuang et al., 2019).

### RNA Sequencing and Analysis

For RNA sequencing, immunoprecipitated RNAs were subjected to quality check and quantification by using the Qubit RNA HS Assay (Thermo Fisher Scientific), and size profiling by using the BioAnalyzer RNA Nano Assay (Agilent). RNA-seq library was constructed by using SMARTer Stranded RNA Kit-Pico Input Mammalian (Takara Bio USA) following the manufacture’s protocol. In brief, 0.4-10 ng of RNA was subjected to heat fragmentation and first-strand cDNA synthesis with SMARTScribe RT enzyme and template-switching oligo (TSO). The cDNA products were then PCR-amplified with simultaneous barcode engineering, and purified by AMPure XP beads (Beckman Coulter). A reaction of the kit’s Control RNA was carried out in parallel. The ribosomal cDNA fragments were depleted using the denatured R-probes of the mammalian kit and the recovery was determined. After AMPure bead purification, the final cDNA libraries were checked by Qubit HS DNA (Thermo Fisher Scientific) and BioA HS DNA Assay (Agilent). The libraries were normalized for effective molar concentrations by quantitative PCR (qPCR) using KAPA Library Quantification Kit Illumina® Platforms (Roche) against the concentration standards. Next-generation sequencing was conducted with SR101nt format (single-end reads, length 101 nt) on a HiSeq 2500 sequencer (Illumina), and obtained 35.4-41.5M reads per sample. The data are strand-specific due to the cDNA orientation anchored at the TSO step. Sample preparation and sequencing work were conducted at the High Throughput Genomics Core of Academia Sinica, Taiwan.

The processed reads were mapped to the human genome (GRCh38) to estimate gene expression level using the Tuxedo protocol (Trapnell et al., 2012). A total of 28,593 expressed genes (FPKM > 0 at least one sample) were detected. NOISeq R package (Tarazona et al., 2015) was used to determine 6,175 differentially expressed genes (DEGs) with probability > 0.9. This set of genes represented Y14-associated transcripts; The correlation coefficient of two duplicate samples was 0.75, indicating substantial reproducibility of the RIP-seq. The pathway enrichment test for identified transcripts was conducted by WebGestalt (Liao et al., 2019) with false discovery rate < 0.05. From the set of DEGs, 349 lncRNAs were identified according to the LNCipedia database (Volders et al., 2019). A total of 104 highly expressed (average FPKM > 10) lncRNAs were selected for further analysis.

### UV-Crosslinking and Immunoprecipitation

HEK293 cells were transiently transfected with an empty vector or FLAG-tagged Y14 (wild-type, SA or WV mutant), eIF4A3 and Ku70. Forty-eight hours post-transfection, cells were UV-crosslinked with 100 mJ/cm^2^ (Stratagene) and lysed in the hypotonic buffer containing 10 mM Tris-HCl (pH 7.5), 10 mM NaCl, 10 mM EDTA, 0.5% Triton X-100 and protease inhibitor cocktail (Roche Applied Science) on ice for 10 min. Subsequently, additional NaCl was added to the lysate to a final concentration of 150 mM. After centrifugation at 13,400 × *g* at 4°C for 15 min, cell debris was removed, followed by immunoprecipitation using anti-FLAG M2 affinity gel (Sigma). After incubation at 4°C for 2 hours, beads were washed with NET-2 buffer. Immunoprecipitates were subjected to immunoblotting and reverse transcription-PCR (RT-PCR) respectively for protein and RNA analysis.

### RNase H Cleavage

HEK293 cells were UV-crosslinked and lysed in hypotonic buffer as described above. After removal of cell debris, anti-FLAG M2 affinity gel and antisense DNA oligonucleotides (5 µM) were added to the lysates and incubated at 4°C for 2 hrs. RNase H (50 units/ml, New England BioLabs) digestion was carried out at 37°C for 1 hour. The beads were subsequently washed with NET-2 buffer and bound proteins were subjected to immunoblotting.

### RNA Pull-down and Mass Spectrometric Analysis

UV-crosslinking and cell lysate preparation were as described above. Biotinylated antisense DNA oligonucleotide probes (100 pmol) and magnetic streptavidin beads (Thermo Fisher Scientific) were added to extracts. After incubation at room temperature for 4 hrs, beads were washed with NET-2 buffer and bound RNAs and associated proteins were subjected to RT-PCR and immunoblotting, respectively. For mass spectrometry, samples were fractionated on SDS-polyacrylamide gels and stained with Coomassie blue. The bands of interest were excised, trypsinized and subjected to nanoAcquity UPLC system (Waters) coupled with the Orbitrap Exploris™ 480 mass spectrometer (Thermo Fisher Scientific).

In brief, peptide mixtures were separated on BEH C18 column (130Å, 1.7 µm, 75 µm x 250 mm, Waters) using a gradient in 30 min from 5% to 35% solvent B (solvent B: 0.1% formic acid in acetonitrile; Solvent A: 0.1% formic acid) at a flow rate of 300 nl/min. The mass spectrometer was operated in data dependent acquisition mode. Full MS resolutions were set to 60,000 at m/z 200 and MS^2^ resolutions were set to 15,000. Isolation width was set at 1.3 m/z. Normalized collision energy was set at 30%. The raw files were searched against an in silico tryptic digest of UniProt human proteome database using the Mascot search engine v.2.6.1 (Matrix Science). The search parameters included the mass tolerance of precursor peptide was set as 10 ppm, and the tolerance for MS/MS fragment was 0.02 Da, cysteine carbamidomethylation as a fixed modification, variable oxidation of methionine and variable deamidation of asparagine or glutamine. Peptide spectrum matches were verified by 1% false discovery rate. We obtained 13529 peptides with a SEQUEST score >20, which represented 3990 proteins in the MASCOT search (Table S3). Among them, 688 proteins showed ≥ 5 protein member hits (Table S4).

### Single-Cell Gel Electrophoresis

This assay was carried out using Comet Assay Kit (Abcam). Briefly, HeLa cells were transiently transfected with siRNAs or GapmeRs. Cells were harvested 48 hrs post-transfection and resuspended at 1× 10^5^ cells/ml in PBS. Cells and comet agarose were mixed at 37°C at 1/10 ratio and 150 μl of the mixture was added onto the comet slide followed by incubation at 4°C for 15 min. The comet slide was immersed in the lysis buffer at 4°C for 1 hour and subsequently in alkaline solution at 4°C for 30 min in the dark. Then the slide was subjected to electrophoresis in TBE running buffer at 20V at 4°C for 30 min. After electrophoresis, the slide was immersed in H_2_O for 2 min followed by fixation with 70% ethanol for 5 min and air dry. Cells were stained with Vista green DNA dye and images were acquired by a laser-scanning confocal microscope (LSM 780, Carl Zeiss). Data processing was performed using OpenComet plugin in ImageJ.

### NHEJ Assays

To assess the NHEJ activity, two systems were adopted. For Cas9-mediated cleavage, the Cas9/sgHPRT expressing vector and double stranded DNA oligonucleotides Ins (25 pmole, Thermo Fisher Scientific) (Du et al., 2018) together with siRNAs, GapmeRs or SMG6-expressing vectors were transfected into HeLa cells. Genomic DNA was collected 48 hrs post-transfection and extracted by PureLink™ Genomic DNA Mini Kit (Thermo Fisher Scientific). For qPCR, the reactions containing 100 ng genomic DNA, specific primers (Table S9) and PerfeCTa SYBR Green FastMix PCR Reagent (Quanta Biosciences) were performed in a LightCycler 480 Real-Time PCR System (Roche). For a GFP-based reporter assay, siRNAs or GapmeRs and the pSCE expression plasmid were transfected into HeLa GFP reporter cells (TopoGEN). Cells were harvested 72 hrs post-transfection. GPF-positive cells were detected by fluorescence-activated cell sorting using 17-color LSR II Analytic Flow Cytometer (BD Biosciences).

### *In Vitro* Pull-down Assay

Recombinant His-tagged Y14 fusion proteins (full-length and ΔC) were described previously (Chuang et al., 2019; Hsu Ia et al., 2005). Recombinant His-tagged human Ku70/Ku80 heterodimer was purchased from Sino Biological. For *in vitro* pull-down assay, 5 μg of His-tagged proteins were incubated with 1 µg of total RNA extracted from HeLa cells and His•Bind Resin (Novagen) at 4°C for 2 hours. After extensive wash, bound RNAs and proteins were respectively analyzed by RT-PCR and immunoblotting.

### Immunoblotting

The procedure of immunoblotting was described previously (Chuang et al., 2019). Antibodies used are listed in Table S7.

### Laser Microirradiation

U2OS cells were seeded in Chambered Coverglass (Thermo Scientific) and transiently transfected with pcDNA-HOTAIRM1-6×MS2 (full-length, ΔE2 or Δ3’) and pcDNA-MCP-GFP or GapmeRs. Laser-microirradiation was performed 48 hrs post-transfection using a laser-scanning confocal microscope (LSM 780, Carl Zeiss) and a 405-nm laser diode. After laser-microirradiation, cells were fixed with 4% paraformaldehyde and permeabilized in 0.5 % Triton X-100 in PBS. Immunofluorescence was performed by sequential incubation with primary and secondary antibodies (Table S7). Nuclei were counterstained in Mounting Medium with DAPI (Sigma). Samples were visualized using a laser-scanning confocal microscope (LSM 780, Carl Zeiss) coupled with an image analysis system.

For live cell imaging, U2OS cells that stably expressed GFP-fused Ku70, Ku80 or MDC1 (Chang et al., 2015) were transiently transfected with pLKO.1-shHOTAIRM1. Laser-microirradiation and time lapse imaging were carried out in a SP5 X inverted confocal microscope (Leica Microsystems) using laser diode 405 nm and 488 nm, respectively.

### Fluorescence *in situ* Hybridization

*HOTAIRM1* was detected by fluorescence *in situ* hybridization (FISH) using bHM1-3 as probe (GENOMICS). After laser-microirradiation, U2OS cells were fixed with 4% paraformaldehyde and permeabilized in 0.5 % Triton X-100 in PBS. After wash with PBS, hybridization was performed in hybridization buffer containing 10 % dextran sulfate, 2×SSC, 10 % formamide, 2 mM ribonucleoside-vanadyl complex and bHM1-3 at 37°C for 24 hours.

After hybridization, cells were wash with PBS and incubated with anti-γH2AX, followed by Texas Red-conjugated anti-Biotin and FITC-conjugated anti-mouse IgG (Table S7). Nuclei were counterstained in Mounting Medium with DAPI. Samples were visualized using a laser-scanning confocal microscope (LSM 780, Carl Zeiss) coupled with an image analysis system.

### Immunofluorescence

For immunofluorescence, U2OS cells after 10 Gy IR were fixed with 4% paraformaldehyde and permeabilized in 0.5 % Triton X-100 in PBS. Cells were incubated with antibodies against γH2AX, Upf1 or SMG6 (Table S7) followed by incubation with FITC-conjugated anti-mouse IgG (Cappel) or Alexa Fluor 568-conjugated anti-rabbit IgG (Thermo Fisher Scientific). Nuclei were counterstained in Mounting Medium with DAPI. To detect the kinetics of foci formation, U2OS cells were irradiated and harvested at indicated time points after IR. Cells were washed with PBS and pre-extracted with CSK buffer containing 0.3 mg/ml RNase A (Brown et al., 2015). After pre-extraction, cells were fixed with 2% PFA and permeabilized with 0.2% Triton X-100 in PBS. Cells were then sequentially incubated with primary antibodies against γH2AX or Ku70 (Table S7) and secondary antibodies. Samples were visualized using a laser-scanning confocal microscope (LSM 780, Carl Zeiss) coupled with an image analysis system.

### Statistical Analysis

Statistical analysis was performed using GraphPad Prism 8.0.

## SUPPLEMENTAL TABLES

**Tables S1-S6, see Excel files**

**Table S1. List of Y14-Associated Transcripts Identified by RIP-seq. Related to Figure 1A**.

**Table S2. Reactome Pathway Enrichment Analysis of Proteins Encoded by Y14-Associated mRNAs. Related to Figure 1B**.

**Table S3. List of *HOTAIRM1*-Associated Proteins Identified by Mass Spectrometry. Related to Figure 6A**.

**Table S4. List of *HOTAIRM1*-Associated Proteins with** ≥ **5 Protein Member Hits in the MASCOT Search. Related to Figure 6B**.

**Table S5. KEGG Pathway Enrichment Analysis of *HOTAIRM1*-Associated Proteins that are Listed in Table S4. Related to Figure 6B**.

**Table S6. List of *HOTAIRM1*-Associated Proteins that Function in mRNA Biogenesis and DNA Damage Repair. Related to Figure 6C**.

**Table S7. List of Antibodies Used for Immunoblotting and Immunofluorescence**

**Table S8. List of siRNAs, Antisense Oligonucleotides, GapmeRs and Biotinylated Oligonucleotides**

**Table S9. List of Primers Used for PCR and RT-PCR**

