## Supplementary figures and images for "lncRNA *HOTAIRM1* Coordinates with RNA Processing Factors in DNA Damage Repair"

### Supplemental Figures

**Supplementary Figure 1**

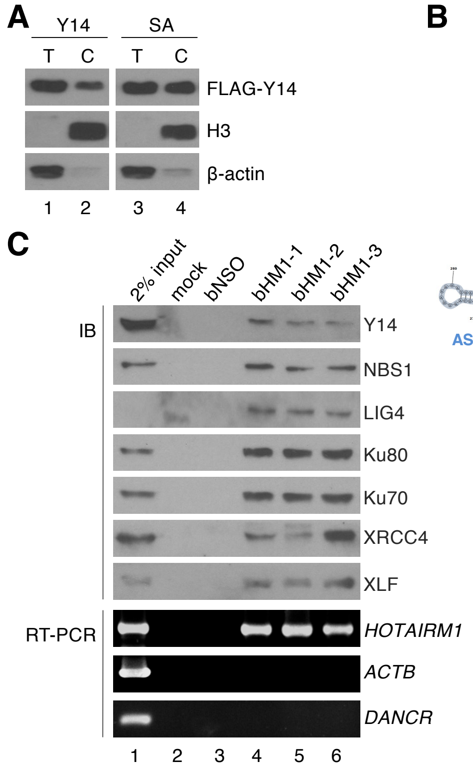


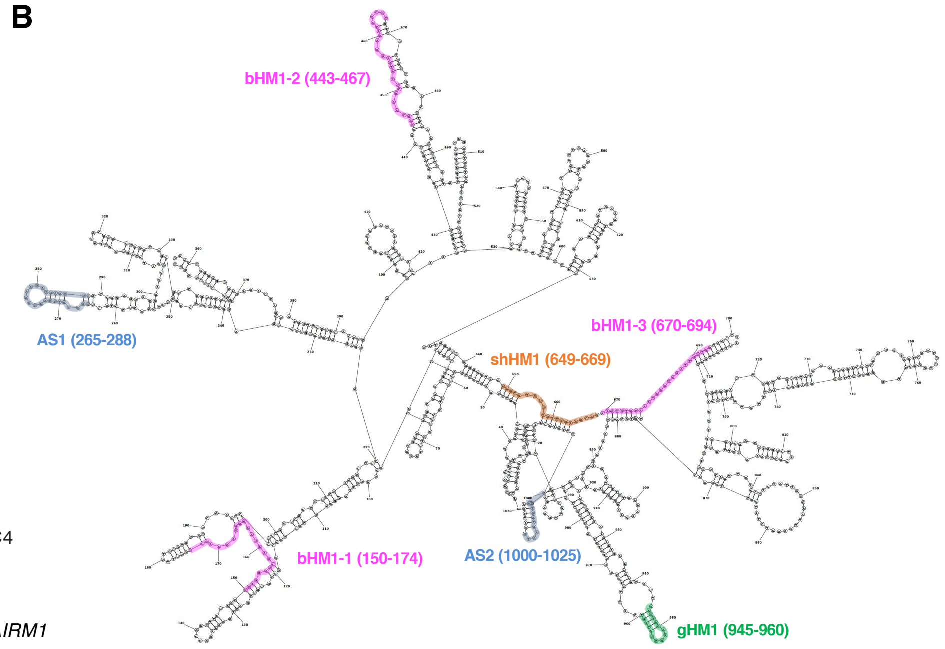

**Supplementary Figure 2**

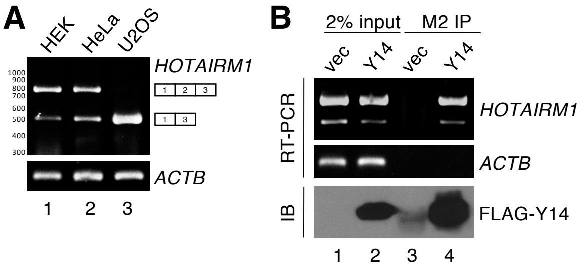

**Supplementary Figure 3**

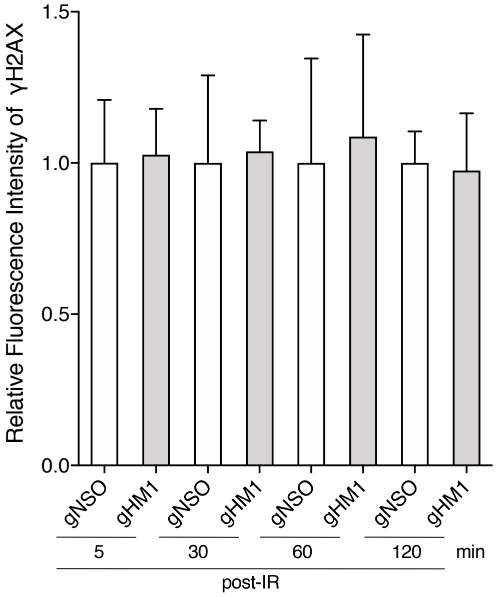

#### Supplementary Figure 4

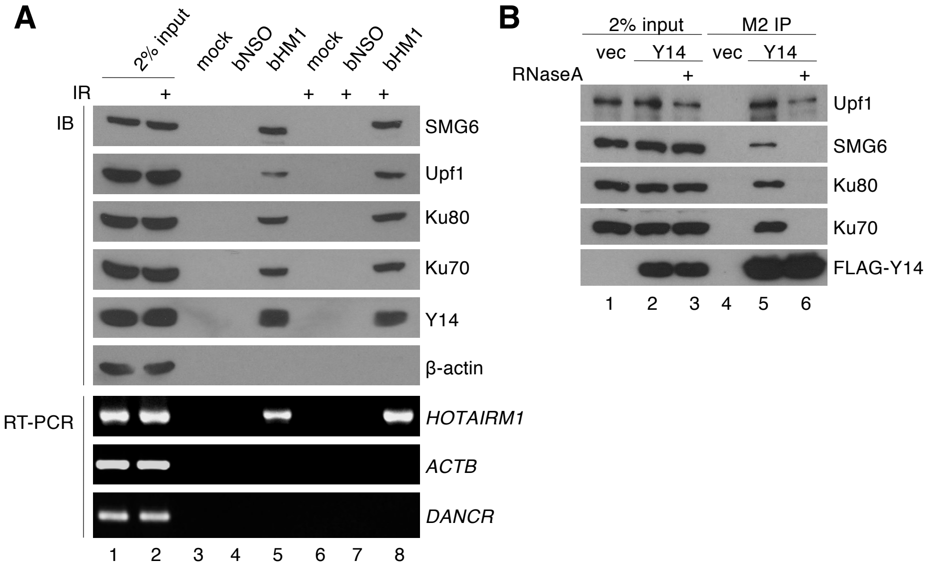
